## Supplemental materials for "The transcriptome of HTLV-1-infected primary cells following reactivation reveals changes to host gene expression central to the proviral life cycle"

### Supplementary materials

#### Supplementary figures and tables

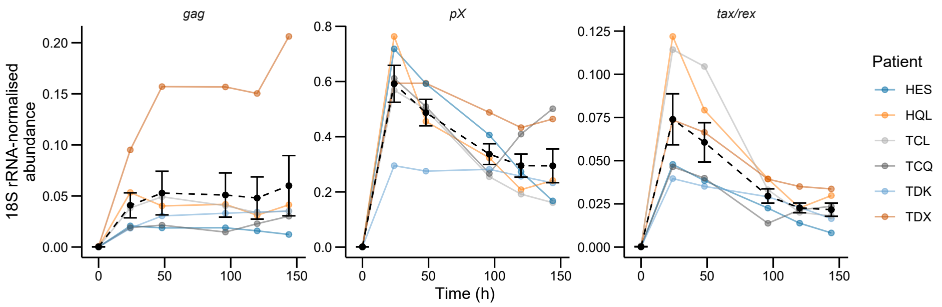


**Figure S1: Proviral transcript expression following reactivation.**
HTLV-1 sense-strand RNA trajectories during *ex vivo* culture, obtained using qRT-PCR. "gag" amplicons correspond to proviral region 2,017–2,203, whilst "pX" corresponds to region 8,000–8,161. *tax/rex* amplicons straddle the second exon junction. Coordinates presented for HTLV-1 sequence AB513134. Dark points and dotted lines represent mean ±SEM.

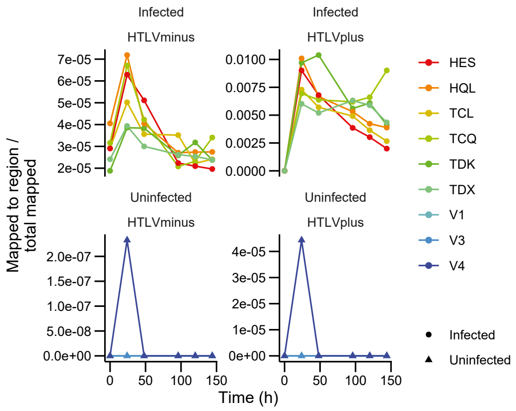


**Figure S2: Aberrant reads mapping to uninfected sample at single timepoint.**
Values represent number reads mapped to HTLV sense or antisense strands normalised to the total number of reads mapped to hg38, with AB513134 appended as an additional chromosome, by STAR.

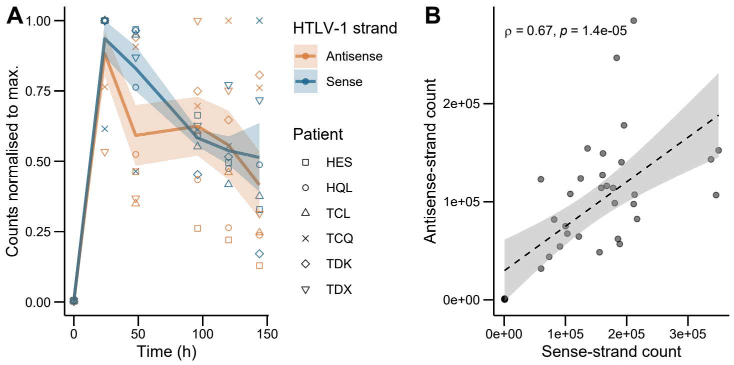


**Figure S3: Antisense-strand transcription correlates with sense-strand reactivation.**
(A) DESeq2-normalised counts for sense and antisense strand transcripts normalised to their maximum values.

(B) DESeq2-normalised counts of reads aligned to the proviral sense or antisense-strand exons. Linear model fit with 95% confidence interval shown. Statistics shown from Spearman's rank correlation test.

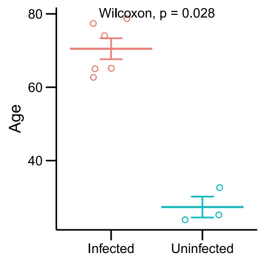


**Figure S4: Age discrepancy between infected and uninfected samples.**
Uninfected controls were not age-matched with infected patients. $p$-value shown from Wilcoxon test.

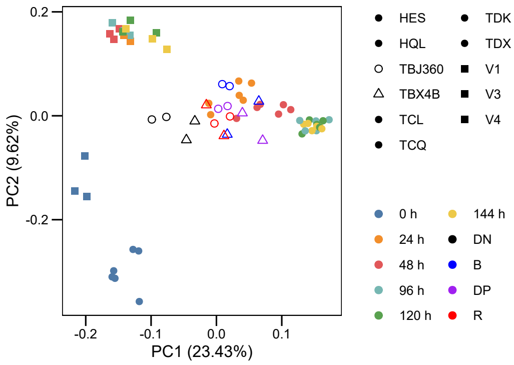


**Figure S5: Clustering of PBMC samples relative to fluorescent timer protein populations described in (1).**
PCA analysis performed on VST-normalised data, subsequently z-scaled to correct for read count discrepancies between datasets.

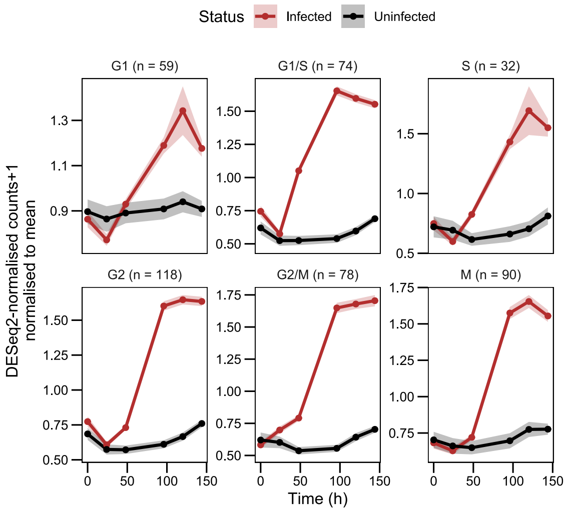


**Figure S6: Cells progress through the cell cycle following the peak in sense-strand expression.**

Mean trajectories of genes with peak expression levels at distinct cell cycle stages. Shaded regions represent ±SEM.

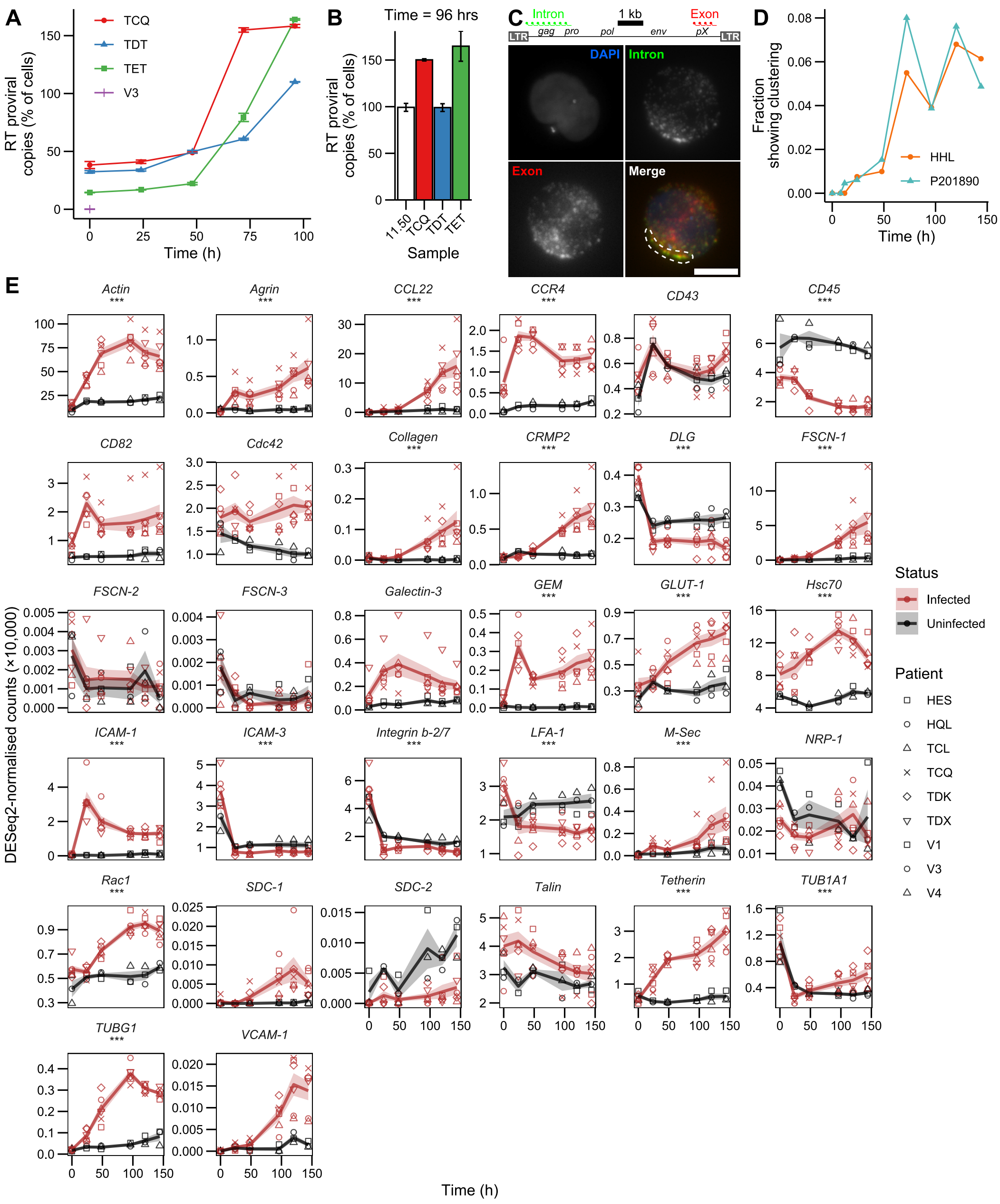


**Figure S7: Changes in genes associated with horizontal infection over expression cycle.** (A) Left: ddPCR PVL measurements for three infected samples and one uninfected control (V3). Right: Repeat measurement of 96-hour infected-cell samples, together with 11.50 positive control for 100% PVL. Error bars represent ±SD of two technical replicates consisting of sample dilutions. (B) Final timepoint from panel A, with 11.50 positive control for 100% PVL included. Error bars ±SEM. (C) Above: Schematic of HTLV-1 provirus and relative positioning of smFISH probes. Below: Example of cell showing clustering of unspliced proviral RNA near cytoplasmic periphery. Scale bar 5 $\mu$m. (D) Quantification of cells with visible clusters of unspliced RNA. (E) Trajectories of genes reported to influence horizontal infection. Asterisks ('***') indicate genes which change significantly over time and relative to uninfected cells. Summary lines and shaded areas represent mean ±SEM.

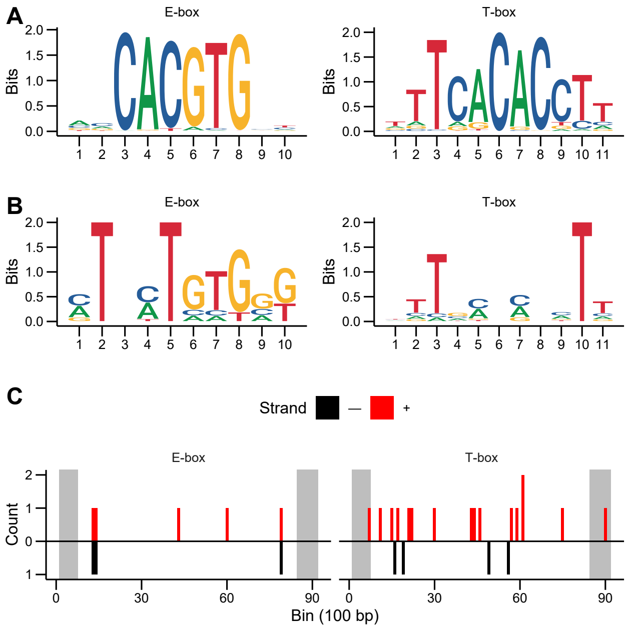


**Figure S8: Distribution of E- and T-box motifs in HTLV-1.**
(A) Sequence logos of E- and T- box motifs obtained from JASPAR (2). (B) Sequence logos of significantly matching ($p < 0.001$) sequences in HTLV. Generated using `ggseqlogo` (3). (C) Distribution of sequences with significant similarity to E-box and T-box motifs along provirus. Grey boxes represent LTRs.

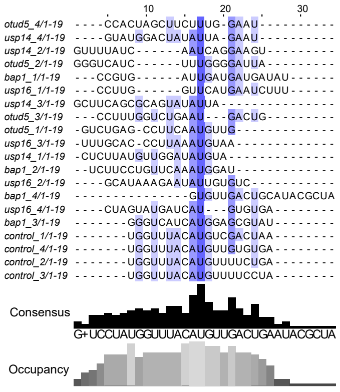


**Figure S9: Alignment of siRNA sequences used for DUB RNAi reveals no overt similarity between co-affected genes.**
Sequences of siRNA fragments used to knockdown DUB transcripts, aligned using Clustal Omega (4).

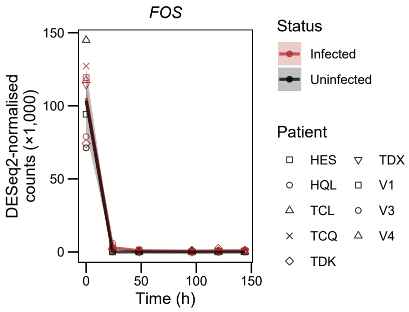


**Figure S10: *FOS* already upregulated at time of 0-hour measurement.**
Lines and shaded areas represent mean ±SEM.

**Table S1: Ranking of histone-associated GO:BP terms from ORA against sense-strand correlates.**

| ID | Description | GeneRatio | BgRatio | BH FDR $p$ |
|---|---|---|---|---|
| GO:0033522 | histone H2A ubiquitination | 11/2555 | 26/16234 | 0.102 |
| GO:0016574 | histone ubiquitination | 16/2555 | 47/16234 | 0.115 |
| GO:0010390 | histone monoubiquitination | 11/2555 | 29/16234 | 0.174 |
| GO:0016570 | histone modification | 87/2555 | 440/16234 | 0.321 |
| GO:0071044 | histone mRNA catabolic process | 5/2555 | 11/16234 | 0.37 |
| GO:0033523 | histone H2B ubiquitination | 5/2555 | 12/16234 | 0.442 |
| GO:0016572 | histone phosphorylation | 11/2555 | 39/16234 | 0.478 |
| GO:0035518 | histone H2A monoubiquitination | 6/2555 | 17/16234 | 0.489 |
| GO:0033127 | regulation of histone phosphorylation | 5/2555 | 13/16234 | 0.489 |
| GO:0031056 | regulation of histone modification | 30/2555 | 139/16234 | 0.489 |
| GO:0031063 | regulation of histone deacetylation | 8/2555 | 27/16234 | 0.535 |
| GO:0070544 | histone H3-K36 demethylation | 4/2555 | 10/16234 | 0.54 |
| GO:0016575 | histone deacetylation | 17/2555 | 76/16234 | 0.604 |
| GO:0031062 | positive regulation of histone methylation | 9/2555 | 35/16234 | 0.626 |
| GO:0034968 | histone lysine methylation | 22/2555 | 106/16234 | 0.661 |
| GO:0016571 | histone methylation | 26/2555 | 129/16234 | 0.663 |
| GO:0033182 | regulation of histone ubiquitination | 4/2555 | 12/16234 | 0.663 |
| GO:0051568 | histone H3-K4 methylation | 12/2555 | 52/16234 | 0.663 |

| | | | | |
|---|---|---|---|---|
| GO:0031058 | positive regulation of histone modification | 18/2555 | 85/16234 | 0.679 |
| GO:0031065 | positive regulation of histone deacetylation | 5/2555 | 17/16234 | 0.679 |
| GO:0080182 | histone H3-K4 trimethylation | 5/2555 | 17/16234 | 0.679 |
| GO:0031060 | regulation of histone methylation | 13/2555 | 60/16234 | 0.715 |
| GO:0008334 | histone mRNA metabolic process | 6/2555 | 24/16234 | 0.75 |
| GO:0070933 | histone H4 deacetylation | 3/2555 | 11/16234 | 0.827 |
| GO:0016578 | histone deubiquitination | 5/2555 | 23/16234 | 0.858 |
| GO:0070076 | histone lysine demethylation | 6/2555 | 29/16234 | 0.877 |
| GO:0070932 | histone H3 deacetylation | 4/2555 | 19/16234 | 0.914 |
| GO:0016577 | histone demethylation | 6/2555 | 31/16234 | 0.922 |
| GO:0051570 | regulation of histone H3-K9 methylation | 4/2555 | 20/16234 | 0.933 |
| GO:0051569 | regulation of histone H3-K4 methylation | 5/2555 | 27/16234 | 0.96 |
| GO:0043984 | histone H4-K16 acetylation | 4/2555 | 21/16234 | 0.96 |
| GO:0010452 | histone H3-K36 methylation | 3/2555 | 15/16234 | 0.96 |
| GO:0051567 | histone H3-K9 methylation | 6/2555 | 34/16234 | 0.97 |
| GO:0043981 | histone H4-K5 acetylation | 3/2555 | 16/16234 | 0.97 |
| GO:0043982 | histone H4-K8 acetylation | 3/2555 | 16/16234 | 0.97 |
| GO:0051571 | positive regulation of histone H3-K4 methylation | 3/2555 | 17/16234 | 0.985 |
| GO:0035404 | histone-serine phosphorylation | 2/2555 | 11/16234 | 0.985 |
| GO:0036124 | histone H3-K9 trimethylation | 2/2555 | 12/16234 | 0.999 |
| GO:0035065 | regulation of histone acetylation | 8/2555 | 54/16234 | 1 |
| GO:0033169 | histone H3-K9 demethylation | 2/2555 | 14/16234 | 1 |
| GO:0035066 | positive regulation of histone acetylation | 4/2555 | 29/16234 | 1 |
| GO:0061647 | histone H3-K9 modification | 6/2555 | 45/16234 | 1 |
| GO:0070734 | histone H3-K27 methylation | 2/2555 | 17/16234 | 1 |
| GO:0016573 | histone acetylation | 20/2555 | 149/16234 | 1 |
| GO:0051573 | negative regulation of histone H3-K9 methylation | 1/2555 | 10/16234 | 1 |
| GO:0031057 | negative regulation of histone modification | 5/2555 | 44/16234 | 1 |
| GO:0034969 | histone arginine methylation | 1/2555 | 11/16234 | 1 |

| | | | | |
|---|---|---|---|---|
| GO:0043967 | histone H4 acetylation | 8/2555 | 68/16234 | 1 |
| GO:0031061 | negative regulation of histone methylation | 2/2555 | 21/16234 | 1 |
| GO:0043486 | histone exchange | 6/2555 | 54/16234 | 1 |
| GO:0090239 | regulation of histone H4 acetylation | 1/2555 | 14/16234 | 1 |
| GO:0043966 | histone H3 acetylation | 5/2555 | 60/16234 | 1 |

**Table S2: Genes included in histone associated terms from ORA of sense-strand correlated genes.**

| ID | Description | BH FDR $p$ | $q$ value | Common name | Entrez ID |
|---|---|---|---|---|---|
| GO:0033522 | histone H2A ubiquitination | 0.102 | 0.0976 | UBR2 | 23304 |
| GO:0033522 | histone H2A ubiquitination | 0.102 | 0.0976 | UBE2A | 7319 |
| GO:0033522 | histone H2A ubiquitination | 0.102 | 0.0976 | KDM2B | 84678 |
| GO:0033522 | histone H2A ubiquitination | 0.102 | 0.0976 | UBE2B | 7320 |
| GO:0033522 | histone H2A ubiquitination | 0.102 | 0.0976 | RNF2 | 6045 |
| GO:0033522 | histone H2A ubiquitination | 0.102 | 0.0976 | TRIP12 | 9320 |
| GO:0033522 | histone H2A ubiquitination | 0.102 | 0.0976 | RYBP | 23429 |
| GO:0033522 | histone H2A ubiquitination | 0.102 | 0.0976 | DTX3L | 151636 |
| GO:0033522 | histone H2A ubiquitination | 0.102 | 0.0976 | BMI1 | 648 |
| GO:0033522 | histone H2A ubiquitination | 0.102 | 0.0976 | BCOR | 54880 |
| GO:0033522 | histone H2A ubiquitination | 0.102 | 0.0976 | RING1 | 6015 |
| GO:0016574 | histone ubiquitination | 0.115 | 0.11 | PAF1 | 54623 |
| GO:0016574 | histone ubiquitination | 0.115 | 0.11 | UBR2 | 23304 |
| GO:0016574 | histone ubiquitination | 0.115 | 0.11 | UBE2A | 7319 |
| GO:0016574 | histone ubiquitination | 0.115 | 0.11 | KDM2B | 84678 |
| GO:0016574 | histone ubiquitination | 0.115 | 0.11 | PARK7 | 11315 |
| GO:0016574 | histone ubiquitination | 0.115 | 0.11 | UBE2B | 7320 |
| GO:0016574 | histone ubiquitination | 0.115 | 0.11 | RNF2 | 6045 |
| GO:0016574 | histone ubiquitination | 0.115 | 0.11 | CDC73 | 79577 |

| | | | | | |
|---|---|---|---|---|---|
| GO:0016574 | histone ubiquitination | 0.115 | 0.11 | TRIP12 | 9320 |
| GO:0016574 | histone ubiquitination | 0.115 | 0.11 | RNF20 | 56254 |
| GO:0016574 | histone ubiquitination | 0.115 | 0.11 | RYBP | 23429 |
| GO:0016574 | histone ubiquitination | 0.115 | 0.11 | DTX3L | 151636 |
| GO:0016574 | histone ubiquitination | 0.115 | 0.11 | BMI1 | 648 |
| GO:0016574 | histone ubiquitination | 0.115 | 0.11 | BCOR | 54880 |
| GO:0016574 | histone ubiquitination | 0.115 | 0.11 | CTR9 | 9646 |
| GO:0016574 | histone ubiquitination | 0.115 | 0.11 | RING1 | 6015 |
| GO:0010390 | histone monoubiquitination | 0.174 | 0.166 | PAF1 | 54623 |
| GO:0010390 | histone monoubiquitination | 0.174 | 0.166 | KDM2B | 84678 |
| GO:0010390 | histone monoubiquitination | 0.174 | 0.166 | RNF2 | 6045 |
| GO:0010390 | histone monoubiquitination | 0.174 | 0.166 | CDC73 | 79577 |
| GO:0010390 | histone monoubiquitination | 0.174 | 0.166 | RNF20 | 56254 |
| GO:0010390 | histone monoubiquitination | 0.174 | 0.166 | RYBP | 23429 |
| GO:0010390 | histone monoubiquitination | 0.174 | 0.166 | DTX3L | 151636 |
| GO:0010390 | histone monoubiquitination | 0.174 | 0.166 | BMI1 | 648 |
| GO:0010390 | histone monoubiquitination | 0.174 | 0.166 | BCOR | 54880 |
| GO:0010390 | histone monoubiquitination | 0.174 | 0.166 | CTR9 | 9646 |
| GO:0010390 | histone monoubiquitination | 0.174 | 0.166 | RING1 | 6015 |

**Table S3: smFISH probe sequences.**

| ID | Sequence (5′ to 3′) | Fluorophore | Source |
|---|---|---|---|
| tax | AGACTCTGTCCAAACCCT | Q670 | (5) |
| tax | CGTAGACTGGGTATCCGA | Q670 | (5) |
| tax | CAGTCGCCTTGTACACAG | Q670 | (5) |
| tax | AACATAGTCCCCCAGAGA | Q670 | (5) |
| tax | CGTGACGATGTAGGCGGG | Q670 | (5) |

| | | | |
|---|---|---|---|
| tax | TGGACAGGTGGCCAGTAG | Q670 | (5) |
| tax | TCCCAGGTGATCTGATGC | Q670 | (5) |
| tax | AACGCGTCCATCGATGGG | Q670 | (5) |
| tax | ACTGTAGAGCTGAGCCGA | Q670 | (5) |
| tax | GGGTCTTAGAGGTTCTCT | Q670 | (5) |
| tax | ATTGGCGGGGTAAGGACC | Q670 | (5) |
| tax | GTTGGGGGTTGTATGAGT | Q670 | (5) |
| tax | GGAGTATTTGCGCATGGC | Q670 | (5) |
| tax | TGGGTTCCATGTATCCAT | Q670 | (5) |
| tax | GACAGGGTTGGGAGGTGC | Q670 | (5) |
| tax | GAGTCCGGGGTCTGGAAA | Q670 | (5) |
| tax | GTGTACAGGTTTTGGGGC | Q670 | (5) |
| tax | ATGCAGACAACGGAGCCT | Q670 | (5) |
| tax | GATGGGGGGGGAAAGCTG | Q670 | (5) |
| tax | TCACATGGGGCAGGAGGG | Q670 | (5) |
| tax | TGGCCGGGGTGGCAAAAA | Q670 | (5) |
| tax | ACATTGGTGAGGAAGGCC | Q670 | (5) |
| tax | CCCCTGTGGTAAGGGAAA | Q670 | (5) |
| tax | ACAGTCCTCGGGTAGAAT | Q670 | (5) |
| tax | GGAAAAGGGTGGTGGGCA | Q670 | (5) |
| tax | CTGTCAGCGTGACGGGTG | Q670 | (5) |
| tax | GAAGGAGGCCGTTTTGCC | Q670 | (5) |
| tax | GTGAGGGTTGAGTGGAAC | Q670 | (5) |
| tax | CCAAATAAGGCCTGGAGT | Q670 | (5) |
| tax | GCGTGCCATCGGTAAATG | Q670 | (5) |

| | | | |
|---|---|---|---|
| tax | CAGGGCCCGGAAATCATA | Q670 | (5) |
| tax | ATGGCTGGCCATCTTTAG | Q670 | (5) |
| tax | GGAGGAGGACTGTAGTAC | Q670 | (5) |
| tax | GGCCTTGGTTTGAAATTT | Q670 | (5) |
| tax | GTAGAAATGAGGGGTGGT | Q670 | (5) |
| tax | ACTGTATGAGGCCGTGTG | Q670 | (5) |
| tax | GGGGATGTTGGTGTATTC | Q670 | (5) |
| tax | GTCATCTGCCTCTTTTTC | Q670 | (5) |
| tax | TTTGGGGCTCATGGTCAT | Q670 | (5) |
| tax | ACTGAGAGGCTCTAAGCC | Q670 | (5) |
| tax | CAGACTTCTGTTTCACGG | Q670 | (5) |
| gag | GGGAAAAGATTTGGCCCAT | Q570 | (5) |
| gag | CGGAATAGGGCTAGCGCTA | Q570 | (5) |
| gag | TTAAGCCAGTGATGAGCGG | Q570 | (5) |
| gag | ATCGTAACTGGAGGGACCG | Q570 | (5) |
| gag | CAGATCCAGACTGGTGTTT | Q570 | (5) |
| gag | GGCTAGGAGGGAGTAGTTA | Q570 | (5) |
| gag | GTATCCTTTTGGGAGTAGG | Q570 | (5) |
| gag | AAATTTCATTCACCCGGCC | Q570 | (5) |
| gag | GCTTGGGTTTGGATGAGTA | Q570 | (5) |
| gag | TTGTGGATCAGAATCCGGG | Q570 | (5) |
| gag | CGTAGGCTCAACATAGGGA | Q570 | (5) |
| gag | GTGGGTGCATGACTGGAAG | Q570 | (5) |
| gag | TGTAGGTCTTTCATTTGCC | Q570 | (5) |
| gag | GGTCTGCATAAACTGGGGG | Q570 | (5) |

| | | | |
|---|---|---|---|
| gag | CAGTGGGGTCAAACTGCTG | Q570 | (5) |
| gag | GAGGTCTTGGAGGTCTTTG | Q570 | (5) |
| gag | ACGAGGGAGGAGCAAAGGT | Q570 | (5) |
| gag | GCCTCTGATATAAGGCTAT | Q570 | (5) |
| gag | CTGTAATACCTCGGGTTTC | Q570 | (5) |
| gag | ACCGGCTAAGGGGTTATAA | Q570 | (5) |
| gag | CCTTGTTGTTGTGGATTGT | Q570 | (5) |
| gag | CAGGAAGGGTCTTTGGCAC | Q570 | (5) |
| gag | CAGGCCTTGGAGGATAGAG | Q570 | (5) |
| gag | TACGAAGGCGTGGTAAGGC | Q570 | (5) |
| gag | AGAGCTATGTTGAGGCGTT | Q570 | (5) |
| gag | TGTTTGCATTGGAGTAGGC | Q570 | (5) |
| gag | CCTAGAGGGCTATTAGTGT | Q570 | (5) |
| gag | TGACAAGCCCGCAACATAT | Q570 | (5) |
| gag | GGGTTTTTTAGGCTGGACA | Q570 | (5) |
| gag | CGGAAGCACGGCTGATTTG | Q570 | (5) |
| gag | TCTTGACATAGGGGGCATG | Q570 | (5) |
| gag | TCTCGCTTCCAGTGAGTTG | Q570 | (5) |
| gag | CTCTGGTTCTGGGATAGTG | Q570 | (5) |
| gag | CAGCGGGGAGGTCTAATAG | Q570 | (5) |
| gag | CCTATGGAGTTTTTTGGGT | Q570 | (5) |
| gag | TGTGGGGGGGGAGGTTAAA | Q570 | (5) |
| gag | GGTTAGGAAGGACTTGCTG | Q570 | (5) |
| gag | GGCAGAATAGATGCTGGGT | Q570 | (5) |
| gag | GGCGGGATCTAACGGTATA | Q570 | (5) |

| | | | |
|---|---|---|---|
| gag | AGCTTCGATAGTCTTTGGG | Q570 | (5) |
| gag | GCTATCGGAAGGACTGTCA | Q570 | (5) |
| gag | CCCCTAATACGGATGTATT | Q570 | (5) |
| gag | AAGTGATCTTGGGTTTGGC | Q570 | (5) |
| gag | AACAATAGGCGTTGTCCGG | Q570 | (5) |
| gag | CCCAGTTGTTTTTGGTATC | Q570 | (5) |
| gag | TTGTAAGGCATCACGACCT | Q570 | (5) |
| gag | CTCAGGGAGGTACAGGACG | Q570 | (5) |
| gag | AGGGAACTGGCTGATTTCG | Q570 | (5) |
